# Non Diaphanous Formin Delphilin Acts as a Barbed End Capping Protein

**DOI:** 10.1101/093104

**Authors:** Priyanka Dutta, Sankar Maiti

**Affiliations:** Department of Biological Sciences, Indian Institute of Science Education and Research-Kolkata Mohanpur - 741246, Nadia, West Bengal

**Keywords:** Formin, Delphilin, Expression, Actin and Barbed end capping

## Abstract

Formins are important for actin polymerization. Delphilin is a unique formin having PDZ domains and FH1, FH2 domains at its N and C terminus respectively. In this study we observed that Delphilin binds to actin filaments, and have negligible actin filament polymerizing activity. Delphilin inhibits actin filament elongation like barbed end capping protein CapZ. *In vitro,* Delphilin stabilized actin filaments by inhibiting actin filament depolymerisation. Therefore, our study demonstrates Delphilin as an actin-filament capping protein.

## INTRODUCTION

Regulated actin dynamics is essential for any organism’s survival. Formins are essential for regulation of actin dynamics. Formins play vital roles as they are key actin nucleator in formation of actin filament structure important for cell functioning. Formins are multi domain proteins, ubiquitously expressed in eukaryotes. Formins are characterized by formin homology-2 (FH2) and formin homology-1 (FH1) domain respectively [1]. FH2 domain nucleates actin in dimeric form, having donut-shaped structure linked by flexible linker [2, 3]. They produce long unbranched linear actin filaments, compared to other nucleators [4].

In lower eukaryotes there are fewer formins, e.g. budding yeast has two formin, Bni1 involved in actin cable and Bnr1 important for cytokinetic ring formation [5]. In fission yeast has three formins For3, cdc12, Fus1; required for cell polarity, cytokinetic ring and fusion of cells respectively [6, 7]. In higher eukaryotes, formins are present in large numbers, like *Arabidopsis* have 21 formins [8]. Mammals have 15 different formins [1]; research has shown that a single cell expresses multiple formins at a time [9, 10]. The specific function of every formin, other than actin nucleation inside the cell are yet to be discovered.

We are interested in the least biochemically characterized formin, called Delphilin. Initially Delphilin discovered as Glutamate δ2 receptor interacting protein [11, 12]. So it was named “Del” from delta and “philin” due its affinity for Glutamate delta2 receptor [9]. Delphilin interacts with Glutamate δ2 receptor, through PDZ domain present at its N-terminal [12]. The importance of this interaction is yet to be elucidated. Delphilin is reported to be selectively expressed in post synaptic nerve terminals of Purkinje cells (PC) [11]. Delphilin mutation causes induction of long term depression (LTD), there are no changes in the morphology of PC but the localization of glutamate δ2 receptor is affected [13].

Role of Delphilin in neuronal plasticity and Delphilin mediated actin cytoskeleton dynamics is poorly understood. Our study attempts to take first steps in filling this void. We have done the biochemical characterization of FH2 domain of Delphilin (Del-FH2). Result shows Delphilin acts as a barbed end capping protein and stabilizes actin filaments *in vitro.*

## RESULTS

### FH2 domain of Delphilin binds F-actin and assembles actin at high concentration

The Del-FH2 or the FH1 and FH2 domains of Delphilin together (Del-FH1FH2) (Fig 1A) were purified as N-terminal 6X His tagged protein (Fig 1B and S1A). To understand the actin binding ability of Del-FH2 we carried out F-actin co-sedimentation assay with purified Del-FH2 and Del-FH1FH2. Our results showed that both Del-FH2 and FH1FH2 fragment coprecipitated in pellet fraction along with F-actin in a concentration dependent manner. Results indicated Del-FH2 was able to bind F-actin as in protein control, Del-FH2 remained in the supernatant (Fig 1C, S1B). Binding to actin filaments confirmed that Del-FH2 behaved similar to FH2 domain of other formins with respect to F-actin binding [14]. Del-FH2 did not have any actin bundling property (Fig S2A).

**Figure 1.**
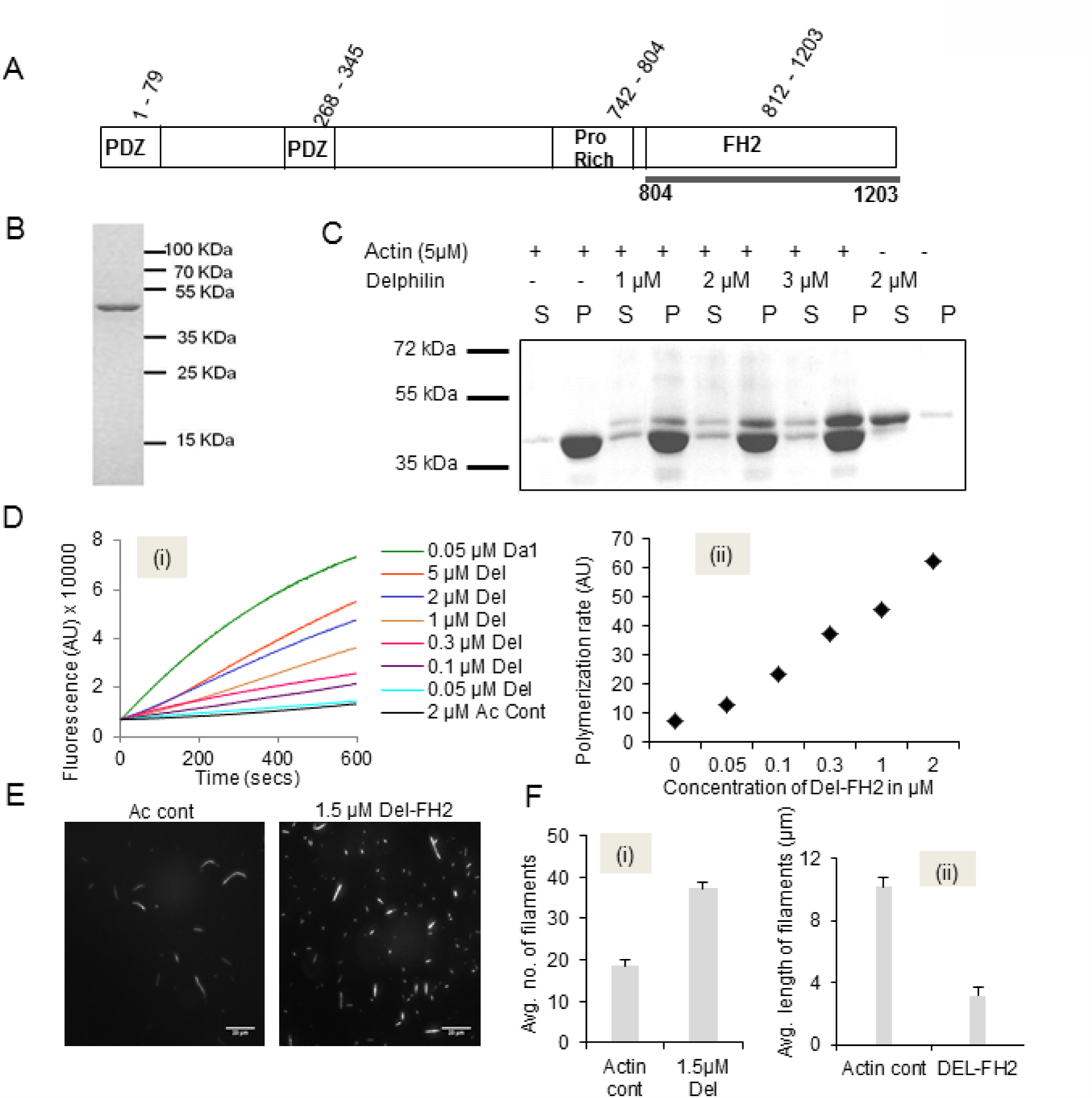
FH2 domain of Delphilin to bind F-actin and assemble actin at slow rate: (A) Schematic illustration of FH1FH2 and FH2 domain of Delphilin. (B) Coomassie stained 12% SDS-PAGE of purified Del-FH2. (C) F-actin co-sedimentation assay of Del-FH2. S and P indicate supernatant and pellet fractions, respectively. (D) (i) Comparison of pyrene-actin nucleation assay with increasing concentration of Del-FH2 and Daam1 (positive control). For *in vitro* kinetics 2μM actin (10% pyrene labelled) was used and indicated (nM and μM) concentration of formin. (ii) Representation of nucleation rate of Del-FH2 concentration. Slopes were taken at 10-100s of time course shown in A. AU, arbitrary units. (E) Fluorescence microscope images of rhodamine phalloidin stained actin filaments in absence and presence of 1.5 μM Delphilin. 0.65 μM rhodamine phalloidin stained actin filaments. Scale bar 20 μm. Magnification is 60X. (F) (i) and (ii) Statistical representation of the number of filaments and the length of the filament (μm) respectively. Error bars represent standard error from 200 filaments for each bar graph.

FH2 domain of formins generally showed *in vitro* actin nucleation activity except Fhod1 [15]. For3 of *Schizosaccharomyces pombe* did not show *in vitro* actin nucleation at low concentration [6]. *In vitro* actin assembly assay of purified Del-FH1FH2/FH2 with 2μM actin monomers (10% pyrene labelled) showed that it was able to assemble actin when present in high micro molar concentrations (Fig 1Di and Fig S1D). However the rate of actin assembly by Del-FH2 was almost negligible than that of positive control Daam1 (Dishevelled associated activator of morphogenesis 1), which was reported to be one of the slowest formin, nucleate actin efficiently [16, 17] (Fig 1Dii). Actin assembly by Daam1 and Delphilin with low salt at 25 nM KCl showed similar pattern of actin assembly at 50 nM KCl (Fig S1C). There was negligible actin nucleation activity of both Daam1 and Delphilin with 0.5 μM of G-actin (Fig S2D).

For further confirmation, actin nucleation by Del-FH2 was observed under fluorescence microscope (Fig 1E). The number of nuclei formed by Delphilin was more than that of actin control (Fig 1Fi) but effective length of the filaments was shorter as compared to actin control (Fig 1Fii). From above data it can be concluded that Del-FH1FH2/FH2 domains form actin nucleus but filament elongation is defective.

In general, Profilin inhibits actin nucleation rate of formins [18]. Del-FH1FH2 mediated actin assembly along with Profilin was almost insignificant and like that of actin control with Profilin (Fig S1Ei). Thus nucleation efficiency of Del-FH1FH2 was found to be irrelevant (Fig S1Eii).

### Inhibition of filament elongation by Delphilin FH1FH2/FH2 domain

FH2 domain of formins involves in the elongation of actin filaments effectively. To see the behaviour of Del-FH2, we had performed actin filament elongation assay accompanying Del-FH1FH2/FH2 and Daam1. Interestingly we had noticed that, Del-FH1FH2/FH2 inhibited of actin filament elongation in a concentration dependent manner (Fig 2A, S3A). Daam1 was able to elongate actin filaments from the barbed end regardless of the concentration (Fig 2B, S3B). When Del-FH1FH2 was present in high concentration, it arrested elongation of actin filaments similar to other barbed end capping proteins (Fig 3Aiv). The microscopic image of filament elongation by Del-FH1FH2 (Fig 3Aiv) was complemented with spectroscopic data of Del-FH2 (Fig 2A, S3A).

Del-FH1FH2/FH2 behaves similarly like plant formin *Arabidopsis* Formin1, inhibiting filament elongation [8]. Del-FH2 act as a barbed end capping protein like Fhod1 [15] and slow down filament elongation in a concentration dependent manner.

Formins FH2 domain has processive capping activity as it captures the filaments tip and able to replace CapZ from filament tips in filament elongation assay. Daam1 was able to replace CapZ from filament tip and carry out elongation of the filament (Fig 2C, S3C). But in case of Delphilin, it did not behave like Daam1 as Delphilin itself inhibit filament elongation (Fig 2D, S3D).

**Figure 2.**
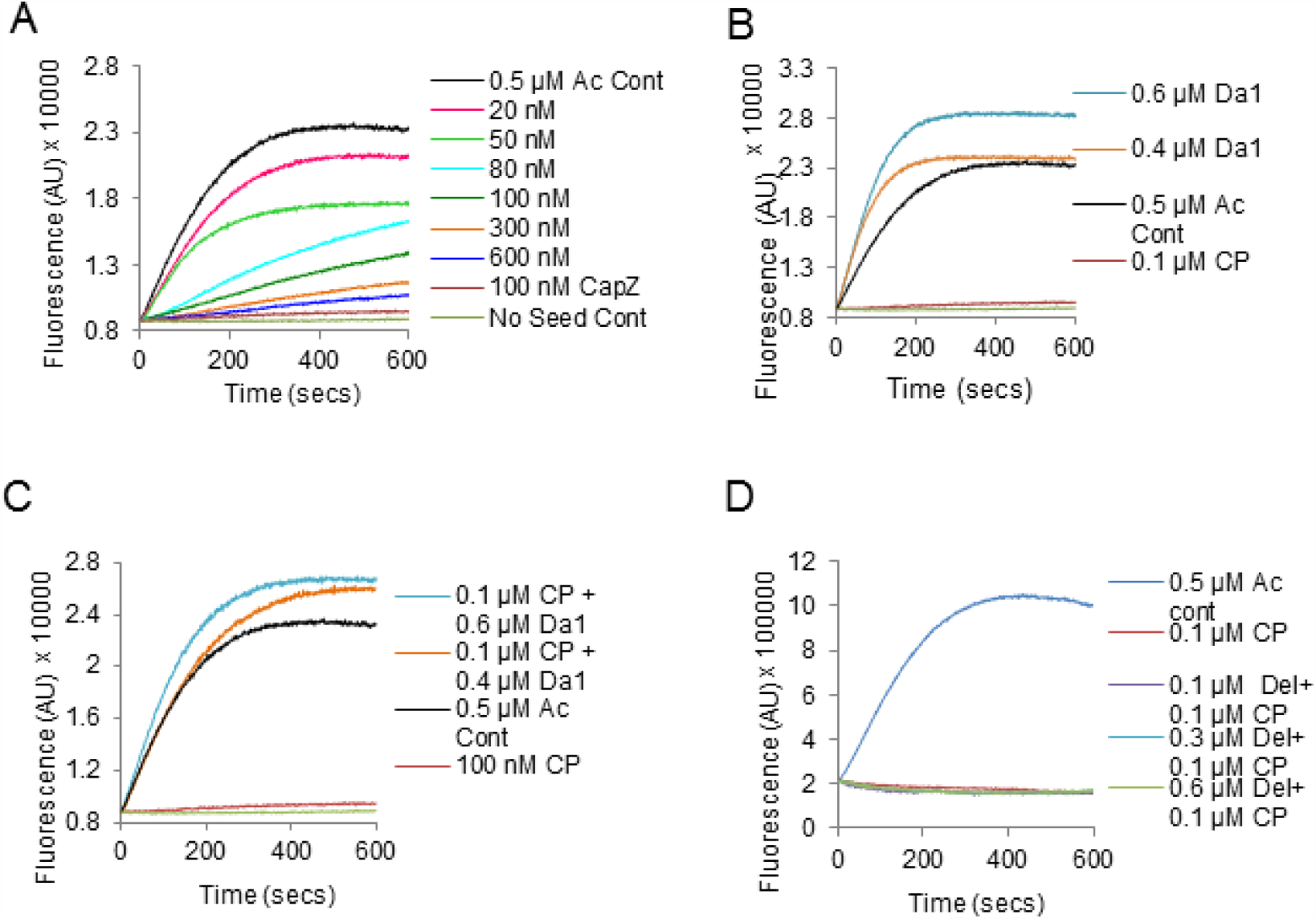
Del-FH2 domain inhibits elongation of actin filaments: (A) Elongation of free barbed end actin filaments along with Del-FH2 in a concentration dependent manner. (B) Free barbed end elongation of actin seeds in presence of Daam1 (Da1). (ii) Plot of elongation rate of Daam1. (C) Daam1 (Da1) (positive control) could easily replace CapZ (CP) at high (nM) concentration. (D) Behaviour of Del-FH2 (Del) in a concentration dependent manner in presence of CP.

It was also observed under fluorescence microscope for filament elongation assay with Daam1 (Fig 3Av) and Del-FH1FH2 (Fig 3Avi) in presence of CapZ and Profilin; Del-FH1FH2 failed to elongate actin filaments. Both spectroscopy and fluorescence microscopy data (Fig S3A) showed inhibition of actin filament elongation by Delphilin and did not show similar pattern like Daam1. Daam1 can replace CapZ in presence of Profilin (Fig 3Av).

**Figure 3.**
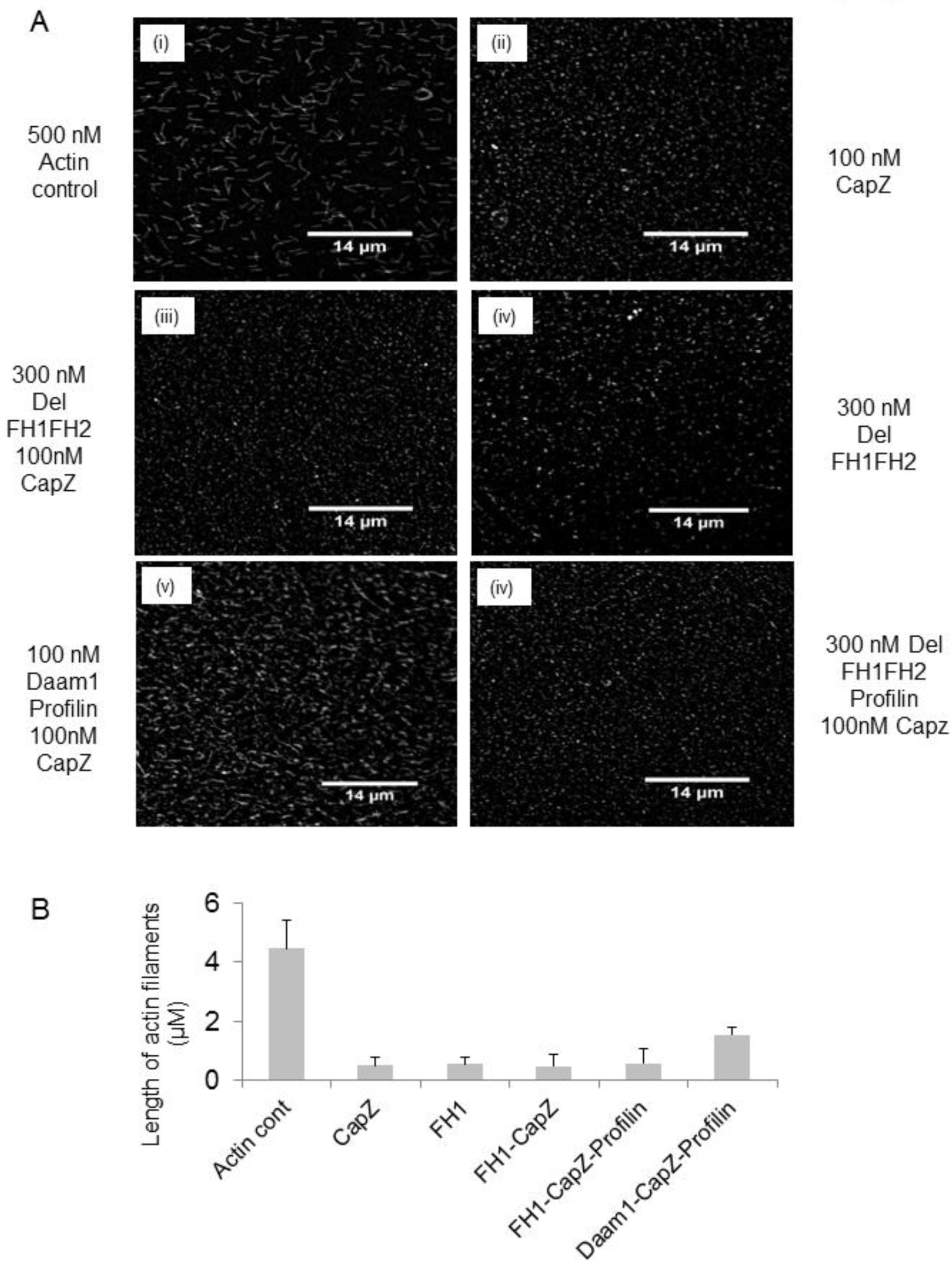
Del-FH2 could not elongate filament in presence of CapZ and inhibit filament elongation: (A) Elongation assay of (i) 0.5 μM actin only, (ii) in presence of CapZ, (iii) in presence of Del-FH1FH2 and CapZ, (iv) in presence of Del-FH1FH2 alone, (v) with CapZ, Profilin and Daam1 and (vi) Del-FH1FH2 plus CapZ and Profilin. Magnifications 60X. Scale bar 14 μm. (B) Bar graphs show inhibition of filament elongation by Del-FH1FH2 domain. Error bars represent standard error from 300 filaments for each bar graph.

For further confirmation actin filament elongation was performed in the presence of 0.4 μM pyrene label RMA and actin seed. Our result showed Del-FH1FH2/FH2 completely inhibits filament elongation but Daam1 did not inhibit filament elongation (Fig S1G, S4Ci, S4Cii). When CapZ and Del-FH1FH2/FH2 are present simultaneously, no filament elongation was recorded from barbed end (Fig S1F S4Bi, S4Bii). Even elongation assay in the presence of 2 μM pyrene label RMA with actin filament seed showed a small increase in fluorescence intensity, which might be due to pointed end growth (Fig S2C).

### Stabilization of actin filaments by Delphilin FH2 domain *in vitro*

*In vitro*, Del-FH2 behaved like a barbed end capping protein. Thus we were interested in actin filament stabilisation activity of Del-FH2 (Fig 4A). Actin filaments *in vitro* depolymerised in a dilution dependent manner. Xenopus cofilin (Xac1) enhanced depolymerisation of actin filaments whereas CapZ stabilized depolymerisation of actin filaments (Fig S2B). The behaviour of Del-FH2 was identical to CapZ (Fig S2B) and stabilized actin filaments in a concentration dependent manner (Fig 4B).

**Figure 4.**
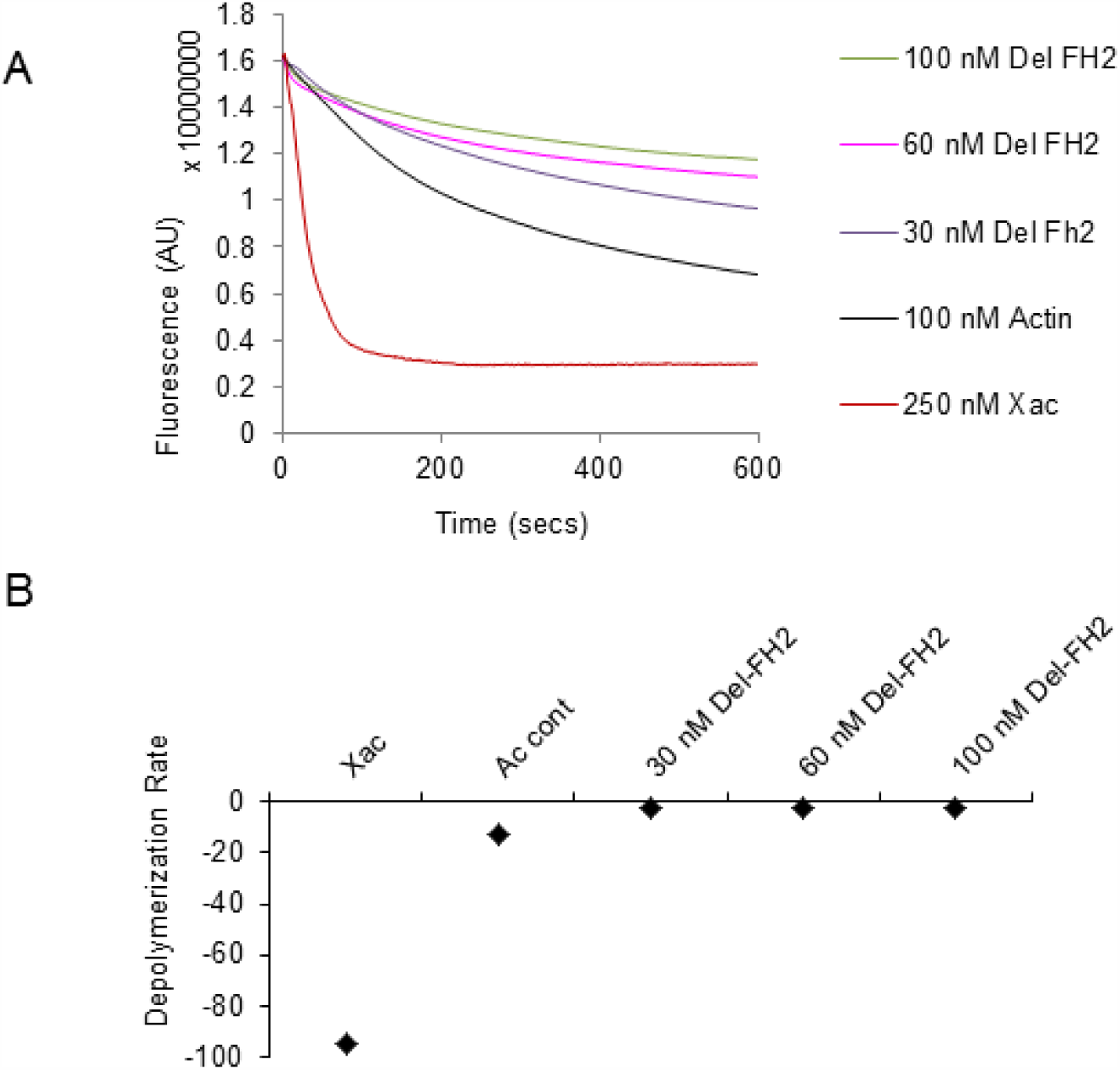
Del-FH2 could stabilize actin filaments in vitro: (A) Del-FH2 could stabilize actin filaments in a concentration dependent manner like barbed end capping proteins. (B) Rate of stabilization of actin filaments by Del-FH2.

## DISCUSSION

In summary, we had characterized Delphilin FH2 as a potent actin capping protein in contrast to other formins which are actin nucleators [1, 15]. Delphilin was reported as glutamate delta2 receptor interacting protein selectively expressed in parallel fibre of postsynaptic cerebellar PC. It has important role in cerebellar LTD, motor coordination and synapse maturation [11]. Delphilin has two forms; L-Delphilin: canonical form and short form S-Delphilin lacking 1st-179^th^ amino acid at the N-terminus [19]. Earlier characterization of N-terminal of Delphilin has shown to interact with glutamate delta2 receptor, while C-terminal consisting of FH1FH2 domain remains unexplored. Our study shows that Del-FH2/FH1FH2 bind to F-actin in a concentration dependent manner (Fig 1C and S1B) like any other formin [1]. But it did not show any actin bundling property (Fig S2A).

Studies on formins from mid 90s to till date have shown that FH2 domain of formins is potent actin nucleators [20]. Research on formins reveals that diaphanous formins can accelerate *in vitro* rate of actin nucleation [18]. We are familiar with the fact that Delphilin is a unique metazoan formin having PDZ domain. Our study also showed that high concentration of Del-FH1FH2/FH2 domain was able to form actin nucleus in a slothful manner i.e. almost negligible compared to Daam1 (Fig 1Dii). But with Profilin, the nucleation capability of Del-FH1FH2 was inconsiderable (Fig S1Ei). The fluorescence microscopy, demonstrated that Del-FH2 increases the number of actin nuclei as compared to actin control (Fig 1E, 1Fi) but size of actin filaments were smaller than actin control (Fig 1Fii). Slow actin assembly by Del-FH2 might be due to some structural differences of FH2 domain of Delphilin, which is yet to be explored.

FH2 domain of formins allows elongation of actin filaments at variable rates except Fhod1 [15]. In case of Daam1; irrespective of the concentration, Daam1 elongates actin filaments (Fig 2Bi, 2Bii) but Delphilin inhibited filament elongation in a concentration dependent manner (Fig 2A, S3A and 3Aiv).

Formins have been reported to alter actin dynamics by protecting filament barbed end from capping proteins [1, 6]. Capping proteins like CapZ [21] bind to barbed end of the filament and stop its elongation. Formins have been reported to replace capping protein i.e. CapZ from barbed end of the filament and allow filament elongation in a stair-stepping manner. Our elongation assay reconfirmed that Del-FH1FH2/FH2 did not perform like Daam1 in presence of CapZ and inhibit elongation of filaments (Fig 2D, 2A, 2C, 3Aiii, S1F and S1G). Our fluorescence microscopy data also showed that Del-FH1FH2 did not function like Daam1 in presence of CapZ and Profilin (Fig 3Av, 3Avi, 3B and S4A). This indicates that Del-FH1FH2/FH2 behaves as a barbed end capping protein and stabilizes actin filaments similar to CapZ (Fig 4A and S2B).

In lower eukaryotes formins are essential for cell polarity but *in vitro* fail to initiate actin assembly at lower concentration [7]. In lower eukaryotes they act as barbed end capping proteins regulated by Profilin [22]. Among metazoan formins, Delphilin was the only formin with PDZ domain at N-terminus and C-terminus with consecutive FH1 and FH2 domains only. From above experiments it can be proposed that Del-FH2 behaves differently when compared with other known metazoan formins as it inhibits filament elongation. Thus, unlike other formins, Del-FH2/FH1FH2 domain acts as a barbed end capping protein.

In future we would like to characterize N-terminal part of Delphilin in order to understand its complex *in vivo* function. Further cellular studies of Delphilin might also shed light on the function of Delphilin in maintaining dendritic spines, spine mobility, learning memory, and synapse stabilization.

## MATERIALS AND METHODS

### Plas mid constructs

Construct of Del-FH2 (804th-1203th) aa was made with forward primer 5’GGCTT**GGATCC**ATGCTATCCCGAGGCGTGG3’ and with reverse primer 5’GCGCC**AAGCTT**TTACCAGGCCAGGGGTGACACC3’ from Delphilin clone (Clone Id: 6841572, Biocat).and Del-FH1FH2 (742nd-1203rd) aa was made by PCR amplification using forward primer 5’GGCCC**GGATCC**TCCGACCACATCCCCCCAC3’ and reverse primer 5’GCGCC**AAGCTT**TTACCAGGCCAGGGGTGACACC3’ respectively and cloned in pET-28a vector (Novagen). Mouse capping protein α1 and β2 subunits were amplified from adult mouse brain cDNA and cloned in pET-28a vector. Human Daam1 (497th-1078th) aa was PCR amplified from human Daam1 clone (Clone Id: 5784628, Thermo Scientific) by using fwd primer 5’GCCGCGGATCCGAGACTACTGAGCATAAGCAAG3’and reverse primer 5’GGAAGGAAAAAAGCGGCCGCTTAGAAATTAAGTTTTGTGATTGGTC3’ and cloned into pet28a vector.

### Protein purification

G-actin was purified from rabbit skeletal muscle acetone powder [23]. Actin was labelled with N-(1-pyrene) iodoacetamide (P-29, Molecular Probes) [24]. Rabbit muscle actin (RMA) and pyrene labelled actin was purified on a Hiprep 16/60 Sephacryl S-200 gel filtration column (GE Healthcare) which was pre equilibrated in G-buffer (2 mM Tris pH 8.0, 0.2 mM ATP, 0.2 mM CaCl2 and 0.2 mM DTT). For purification of proteins; expression constructs were transformed into *Escherichia coli* BL21 DE3 (Stratagene). Cells were cultured in rich media at 37°C until OD_600_ was 0.5-0.6 (20 gm Tryptone/liter, 10 gm Yeast Extract/liter, 0.58 gm NaCl/liter, 0.18 gm KCl/liter and 30 μg/ml Kanamycin). Protein expression was induced with 0.5 mM IPTG and cultures were further grown at 19°C O/N. After 14 hrs, cells were harvested, and stored at −80°C [25].

All steps for protein purification were done at 4°C or on ice [17]. Frozen cell pellets were taken and re-suspended in lysis buffer (50 mM Tris-Cl pH 8.0, 100 mM NaCl, 30 mM Imidazole pH 8.0, 1 mM DTT(USB), IGEPAL(Sigma-aldrich), Thesit (Sigma-Aldrich), Protease inhibitor cocktail (1mM Phenylmethylsulfonyl fluoride, Benzamidine hydrochloride, Leupeptin, Aprotinin, PepstatinA) and lysed by sonication. It was then centrifuged at 12000 rpm for 15 minutes. Supernatant was incubated with 50% Ni-NTA slurry of the beads (Qiagen) for 4 hours then washed thrice with wash buffer (50 mM Tris-Cl pH 8.0, 20 mM NaCl, 30 mM Imidazole pH 8.0). Protein was eluted with elution buffer (50 mM Tris-Cl pH8.0, 20 mM NaCl, 250 mM Imidazole pH8.0 and 5% glycerol). Purified protein was dialyzed overnight in HEKG_5_ buffer (20 mM HEPES, 1 mM EDTA, 50 mM KCl and 5% Glycerol) overnight and further stored in HEKG_10_ buffer. Expression and purification of Daam1, mouse capping protein α1 and β2 construct, was carried out following same protocol except during purification of capping protein α1 and β2 resuspended cells were mixed during sonication.

Xenopus cofilin in pGEX (GST-Xac1) vector was a kind gift from Prof JR Bamburg [26]. GST-tagged Xac1 in *E. coli* was expressed at 37°C with IPTG induction for 4 hours. Harvested cells were resuspended in lysis buffer (50 mM Tris-Cl pH 8.0, 150 mM NaCl, 1 mM DTT, 1 mM EDTA, 0.5% IGEPAL with Protease inhibitor cocktail) then lysed by sonication and purified on a GST matrix (Thermo Scientific). Protein was eluted with reduced glutathione.

### Actin Filament Co-sedimentation Assay

Actin was polymerized for one and half hour at 25°C in F-buffer (10 mM Tris-Cl pH8.0, 0.2 mM DTT, 0.7 mM ATP, 50 mM KCl, 2 mM MgCl2). Different concentrations of Del-FH2 were added and incubated for 15 minutes. Concentration of actin was 5μM in each case with one actin and protein control only. After completion of incubation, samples were centrifuged at 310 x 1000 g for 45 minutes in TLA-100 rotor (Beckman Coulter). Subsequently supernatant was removed and mixed with SDS-PAGE sample loading buffer, while pellet fraction was reuspended in F-buffer and SDS-PAGE sample loading buffer. Samples were then analysed on 12% SDS PAGE and coomassie staining. For bundling experiment similar reaction was used and centrifuged at low speed 9.2 x 1000 g for 10 minutes [20]. These experiments were done thrice.

### Actin nucleation assay by Fluorescence Spectroscopy

Both rabbit muscle actin (RMA) and pyrene actin were mixed to prepare 12 μM 10% pyrene actin in G-Buffer. This mixture was incubated for 2 minutes at room temperature with MgCl_2_ and EGTA. Different concentrations of the desired protein were added and the volume was adjusted with G-buffer and HEKG5. Just prior to polymerization reaction, 20X Initiation Mix (1 M KCl, 40 mM MgCl2, 10 mM ATP) was added. Fluorescence was measured at 365 nm (excitation) and 407 nm (emission) wavelength at 25°C in fluorescence spectrophotometer (QM40, Photon Technology International, Lawrenceville, NJ) [22]. Actin nucleation assay was repeated five times.

### Fluorescence microscopy of phalloidin stained actin filaments

Actin filaments stained with rhodamine phalloidin were seen *in vitro* under fluorescence microscope [27]. Actin alone, and with Delphilin was polymerized with 20X initiation mix. Reaction mixture was diluted with Antifade reagent (Invitrogen). A diluted sample of 10 μl was applied to grease free slide and covered with poly-D-lysine (Sigma Aldrich) coated cover slips (Blue Star 12 mm diameter). Filaments were observed after 20 minutes under fluorescence microscope (Nikon Motorized Inverted Research Microscope, Ti-E equipped with 60X, 1.2-numerical aperture Plano Apo objective) and the images were taken utilising (EvolveTM – 512 Delta CCD) camera. The same procedure was carried out for elongation assay. Actin filaments were sheared with 27 gauge needle for 5 passages and images were taken. Fluorescence microscopy was done twice to observe the change in the length of actin filaments. Number and length of the filaments were counted using ImageJ software. Data was collected, entered and analysed using the Microsoft Excel 2007 statistical analysis tools.

### Filamentous actin elongation assay by Fluorescence Spectroscopy

Unlabeled G-actin was allowed to form F-actin with F-buffer and 10X IM mix (20 mM MgCl2, 5 mM ATP, 500 mM KCl) for 50 minutes at 25°C. 10 μM of 10% pyrene labeled actin stock was prepared in G-Buffer. F actin was sheared by passing through 27 gauge needle 5 times. Immediately 20 μl of this solution was added to mixture of G-buffer and 10% pyrene labelled actin. 10X IM mix was added to it and time-based fluorescence acquisition was initiated. The fluorescence spectroscopy parameters were same as nucleation assay [22]. This experiment was repeated five times to check the abnormal behaviour of Del-FH2.

### Filament stabilization assay by Fluorescence Spectroscopy

7 μM G-actin with 50% pyrene labelled were polymerized in F-buffer (10 mM Tris-Cl pH8.0, 0.2 mM DTT, 0.7 mM ATP, 50 mM KCl, 2 mM MgCl2). Time course of depolymerisation of 7 μM actin filaments after dilution to 0.1 μM were recorded in presence of Xac, various concentration of CapZ and Del-FH2. Fluorescence spectroscopy parameters were same as previous experiments. Stabilization assay was done in duplicate.

## Author’s Contribution

SM and PD conceived and designed experiment. Experiments were performed by PD. Data analyzed: SM and PD. Paper writing: SM and PD.

## Ethical consideration

All experiment protocols were approved by Institutional Animal Ethical Committee of Indian Institute of Science Education and Research-Kolkata. Animals were handled with care and according to the rules of Committee for the Purpose of Control and Supervision of Experiments on Animals (CPCSEA, Govt. of India). CPCSEA registration number of Institute is 1385/ac/10/CPCSEA.

## Acknowledgement

PD thanks Indian Institute of Science Education and Research-Kolkata, India for providing her fellowship. GST-XAC was a kind gift from Prof. JR Bamburg to SM. This work was supported by grants from Department of Biotechnology, India (BT/PR14369/Med/30/526/2010 dated 31/10/2011), Council of Scientific and Industrial Research, India (37(1500)/11/EMR-II, dated 22/12/2011).

---

**Figure S1.**
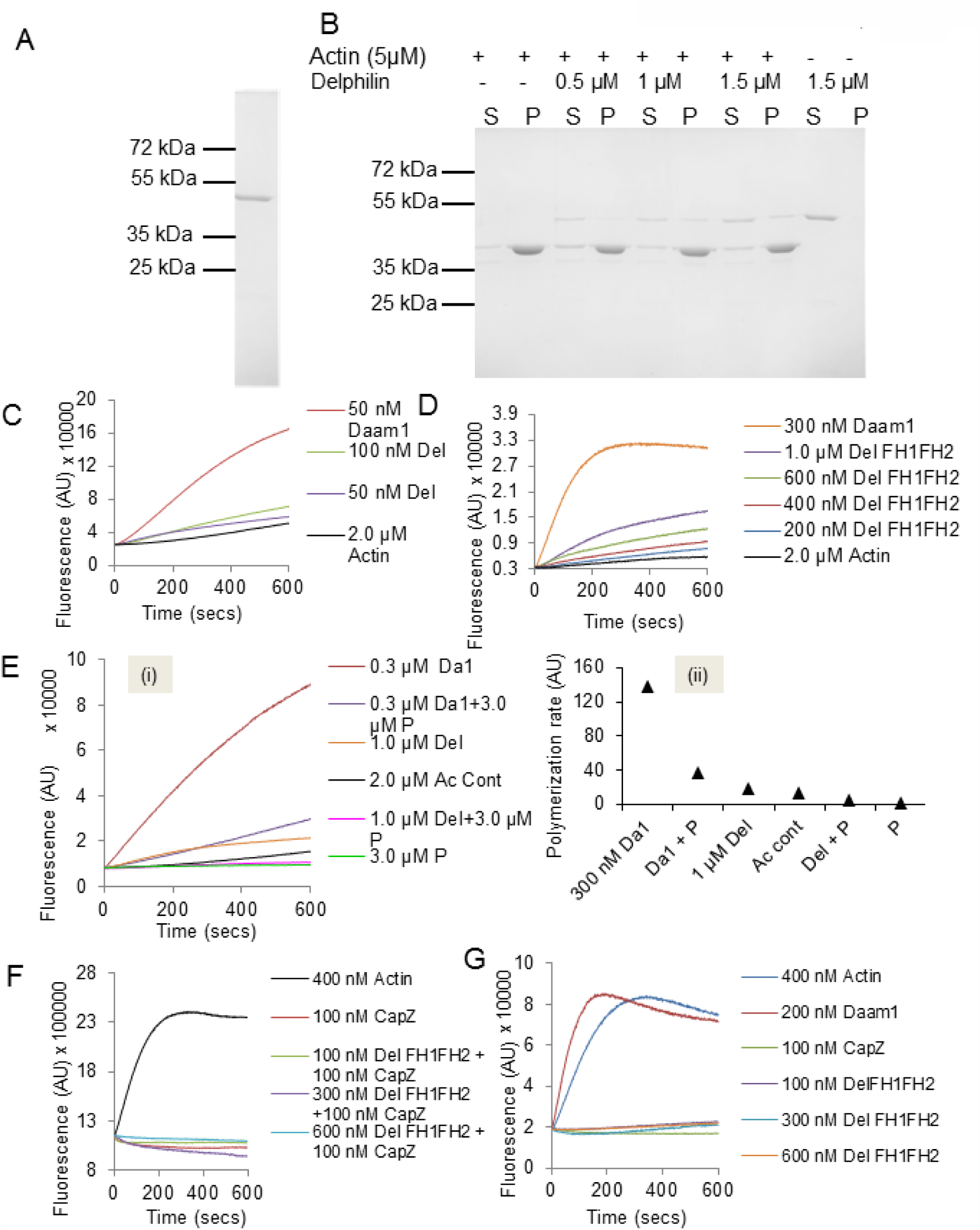
Del-FH1FH2/FH2 domain does not behave as other metazoan formins under different conditions: (A) Coomassie stained 12% SDS-PAGE of Del-FH1FH2. (B) F-actin co-sedimentation assay of Del-FH1FH2. S and P indicate supernatant and pellet respectively. (C) In presence of 25 mM KCl Del-FH2 domain did not behaves like that of metazoan formin Daam1. (D) The ability to nucleate actin filaments by Del-FH1FH2 is negligible compared to Daam1. (E) Nucleation efficiency of Del-FH1FH2 (Del) and Daam1 (Da1) along with Profilin (P). (ii) Comparison of rate of nucleation in presence of Profilin. (F) In presence of 0.4 μM actin seeds Del-FH1FH2 did not replace CapZ. (G) Along with 0.4 μM actin seeds Del-FH1FH2 inhibit filament elongation.

**Figure S2.**
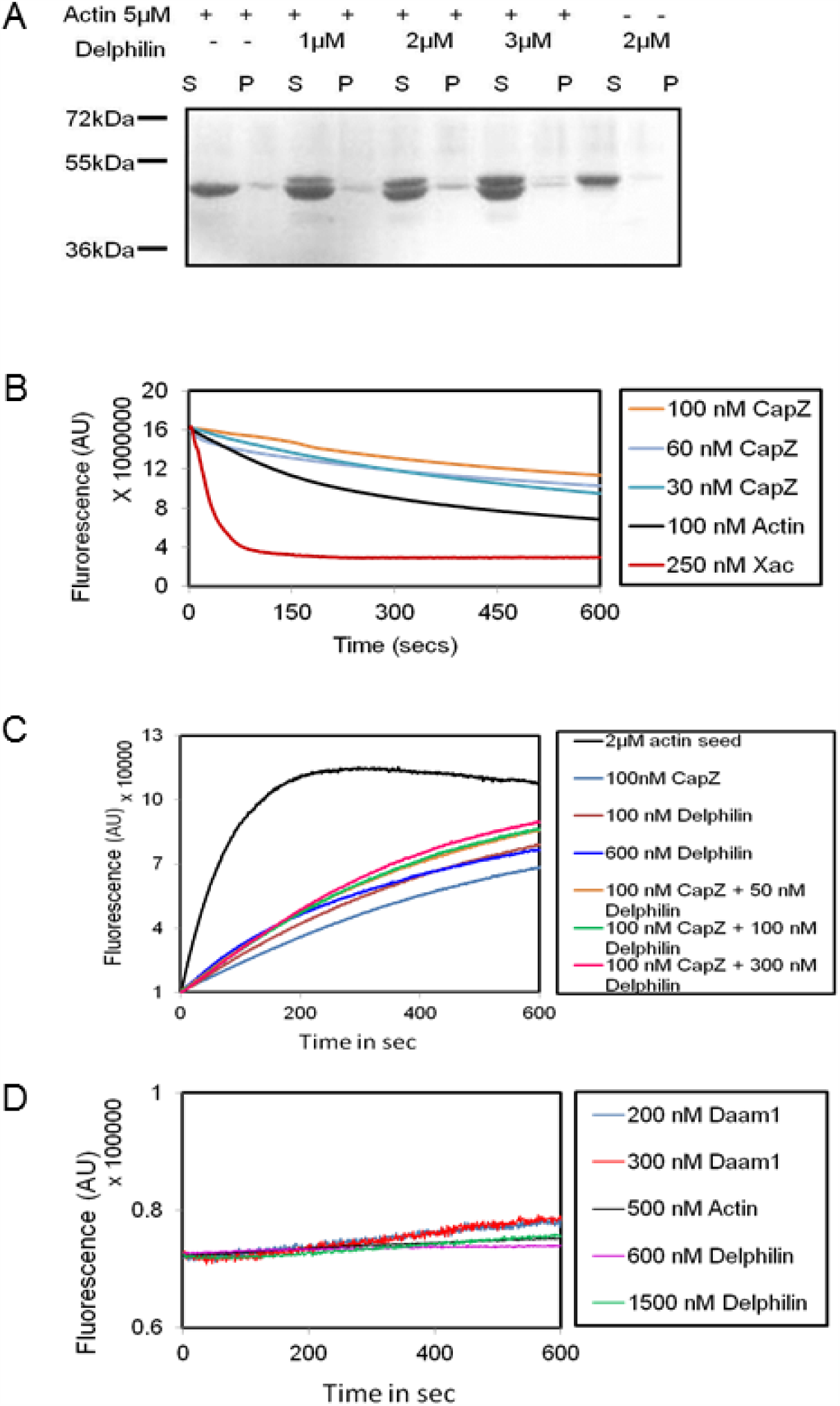
(A) F-actin low speed co-sedimentation assay of Del-FH2. S and P indicate supernatant and pellet respectively. (B) Stabilization of actin filaments by CapZ. (C) Filament elongation assay of Delphilin with 2 μM G-actin concentration. (D) Nucleation assay of Delphilin and Daam1 with 500 nM G-actin.

**Figure S3.**
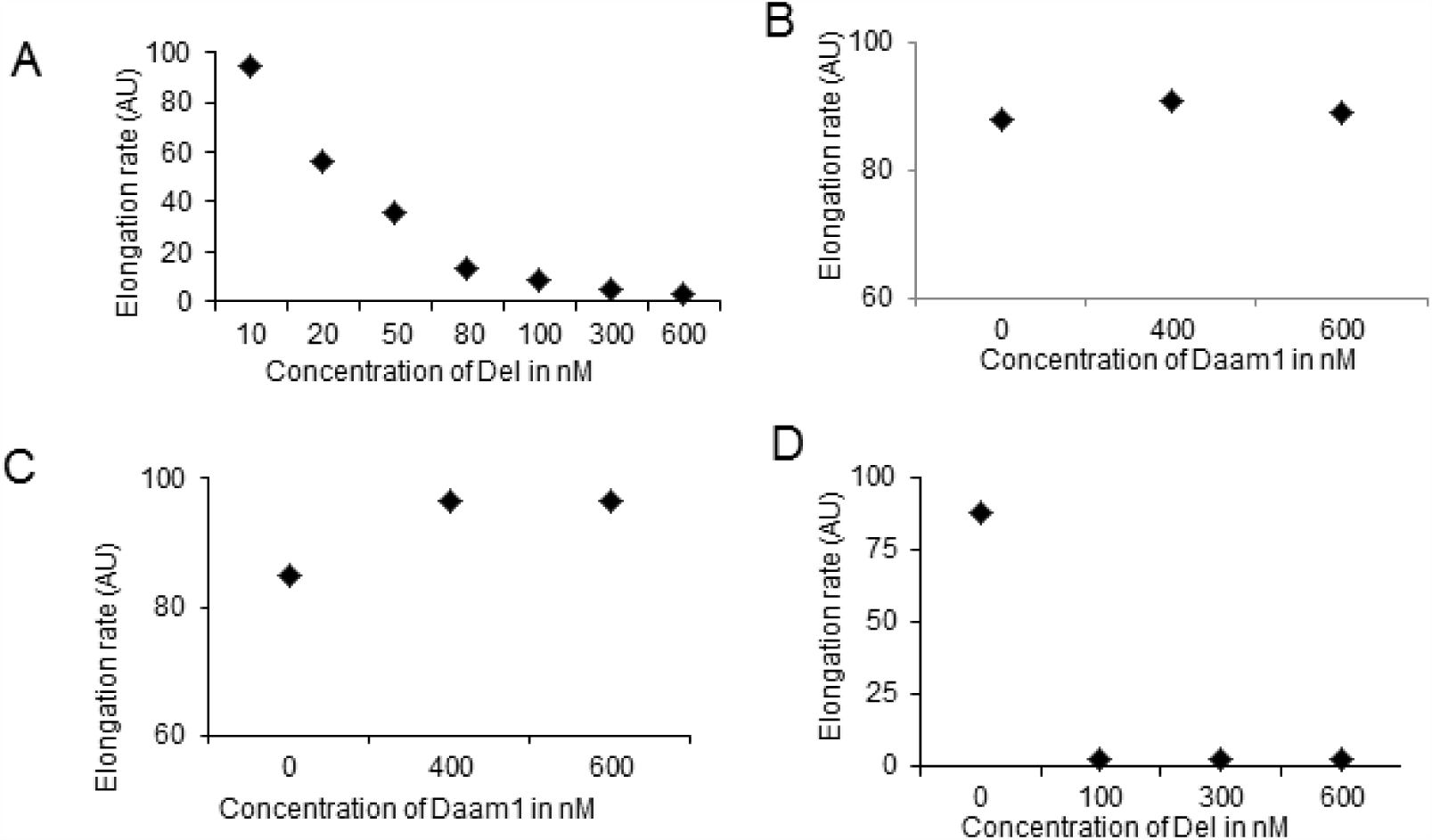
Rate calculation of inhibition of filament elongation: (A) Rate of elongation by Del-FH2 as a function of concentration. Slopes were taken at 10-100s of time course shown in 2A. (B) Plot of elongation rate of Daam1. (C) Rate of replacement of CapZ by Daam1 (Da1). (D) Rate of filament elongation by Del in presence of CapZ (CP).

**Figure S4.**
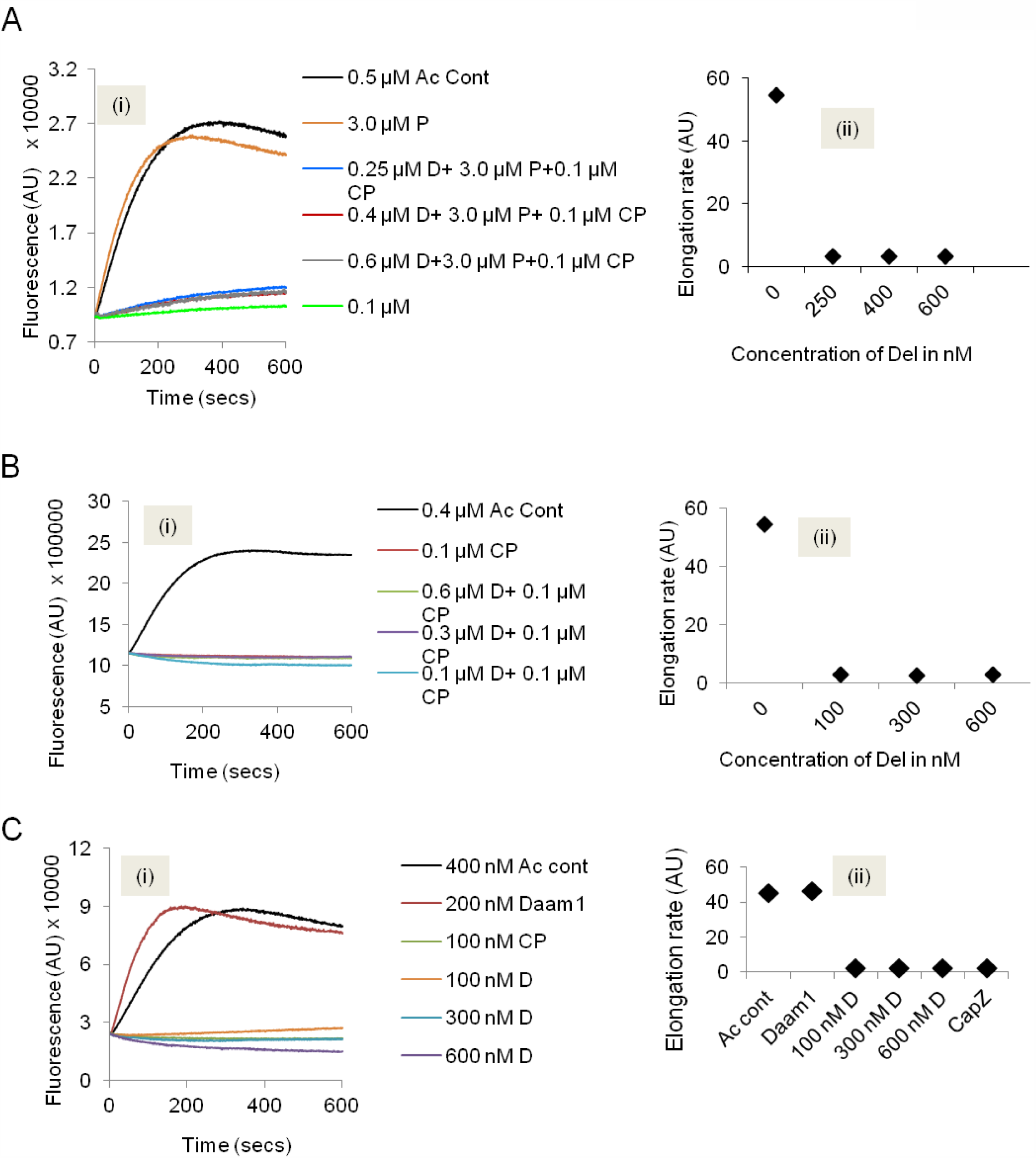
Delphilin could not elongate actin filament from barbed end in presence of CapZ: (A) (i) Activity of Del-FH1FH2 (D) in a concentration dependent manner along with Profilin (P) and CapZ (CP). (ii) Rate of elongation of Delphilin with Profilin and CapZ. (B) (i) Along with 0.4 μM actin seed Delphilin inhibits actin filament elongation. 0.4 μM actin seeds were used to see barbed end kinetics only. (ii) Rate of actin filaments elongation from barbed end at 0.4 μM. (C) (i) Filament elongation from barbed end by Delphilin in presence of 0.4 μM actin seeds and CapZ. (ii) Rate of elongation of Delphilin with CapZ (CP).

